# Inositol acylation of phosphatidylinositol mannosides: a rapid mass response to membrane fluidization in mycobacteria

**DOI:** 10.1101/2022.07.23.501223

**Authors:** Peter P. Nguyen, Takehiro Kado, Malavika Prithviraj, M. Sloan Siegrist, Yasu S. Morita

## Abstract

Mycobacteria share an unusually complex, multilayered cell envelope, which contributes to adaptation to changing environments. The plasma membrane is the deepest layer of the cell envelope and acts as the final permeability barrier against outside molecules. There is an obvious need to maintain the plasma membrane integrity, but the adaptive responses of plasma membrane to stress exposure remain poorly understood. Using chemical treatment and heat stress to fluidize the membrane, we show here that phosphatidylinositol (PI)-anchored plasma membrane glycolipids known as PI mannosides (PIMs) rapidly remodel their structures upon membrane fluidization in *Mycobacterium smegmatis*. Without membrane stress, PIMs are predominantly in a tri-acylated form: two acyl chains of PI moiety plus one acyl chain modified at one of the mannose residues. Upon membrane fluidization, the fourth fatty acid is added to the inositol moiety of PIMs, making them tetra-acylated variants. PIM inositol acylation is a rapid response independent of *de novo* protein synthesis, representing one of the fastest mass conversions of lipid molecules found in nature. Strikingly, we found that *M. smegmatis* is more resistant to the bactericidal effect of a cationic detergent after benzyl alcohol preexposure. We further demonstrate that fluidization-induced PIM inositol acylation is conserved in pathogens such as *Mycobacterium tuberculosis* and *Mycobacterium abscessus*. Our results demonstrate that mycobacteria possess a mechanism to sense plasma membrane fluidity change. We suggest that inositol acylation of PIMs is a novel membrane stress response that enables mycobacterial cells to resist membrane fluidization.

## Introduction

The success of *Mycobacterium* species as human pathogens is partly due to its complex and multilayered cell envelope. It comprises an outer membrane (mycomembrane), a peptidoglycan-arabinogalactan cell wall, a periplasmic space, and a plasma membrane (1–4). The plasma membrane is a typical phospholipid bilayer with phosphatidylethanolamine and cardiolipin as major components. Unlike many other bacteria, however, mycobacteria also produce phosphatidylinositol (PI) as a major phospholipid (5). Reminiscent of glycosyl phosphatidylinositols (GPIs) in eukaryotes, PI is further modified by carbohydrates.

One class of such glycosylated PIs is PI mannosides (PIMs), which are uniquely found in mycobacteria and related bacteria in the Actinobacteria phylum. PIMs may carry up to six mannoses and four fatty acyl chains and are believed to be constituents of the plasma membrane (6). However, earlier studies suggest that PIMs may also be exposed on the cell surface (7, 8) and the fact that multiple host immune receptors, such as the macrophage mannose receptor, DC-SIGN, and DCAR, recognize PIM species further support that at least a fraction of these glycolipids may be exposed on the cell surface (9–11).

PIM biosynthesis takes place in the plasma membrane. In the first step, the mannosyltransferase PimA transfers a mannose residue from GDP-mannose to the 2-OH of the inositol ring of PI, forming PIM1 (Fig. 1) (12). The second step is likely the transfer of the second mannose to the 6-OH of the inositol, mediated by PimB’ (13, 14). The acyltransferase PatA then transfers a palmitoyl moiety (acyl chain) to the 6-OH of mannose, which is linked to the 2-OH of the inositol, forming mono-acylated PIM (AcPIM2) (15) (Fig. 1). An additional palmitoyl moiety may be added to the 3-OH of the inositol to form diacylated PIM (Ac_2_PIM2), but the enzyme that mediates this reaction remains to be identified. AcPIM2 is further modified by additional four mannoses to become AcPIM6, and similar to AcPIM2, it may be acylated to become Ac_2_PIM6. AcPIM2, Ac_2_PIM2, AcPIM6, and Ac_2_PIM6 are the predominant forms of PIMs, accumulating in the cell envelope (Fig. 1). PimE is the only mannosyltransferase with an established role in the biosynthesis of AcPIM6 from AcPIM2 (16). As a polyprenol-phosphate-mannose-dependent α1,2 mannosyltransferase, PimE mediates the synthesis of AcPIM5 from AcPIM4, committing the pathway to AcPIM6 synthesis. AcPIM4 is a key branch-point intermediate, from which an unknown α1,6 mannosyltransferase can take the biosynthetic pathway into another direction to produce larger lipoglycans such as lipomannan (LM) and lipoarabinomannan (LAM) (17).

**Figure 1.**
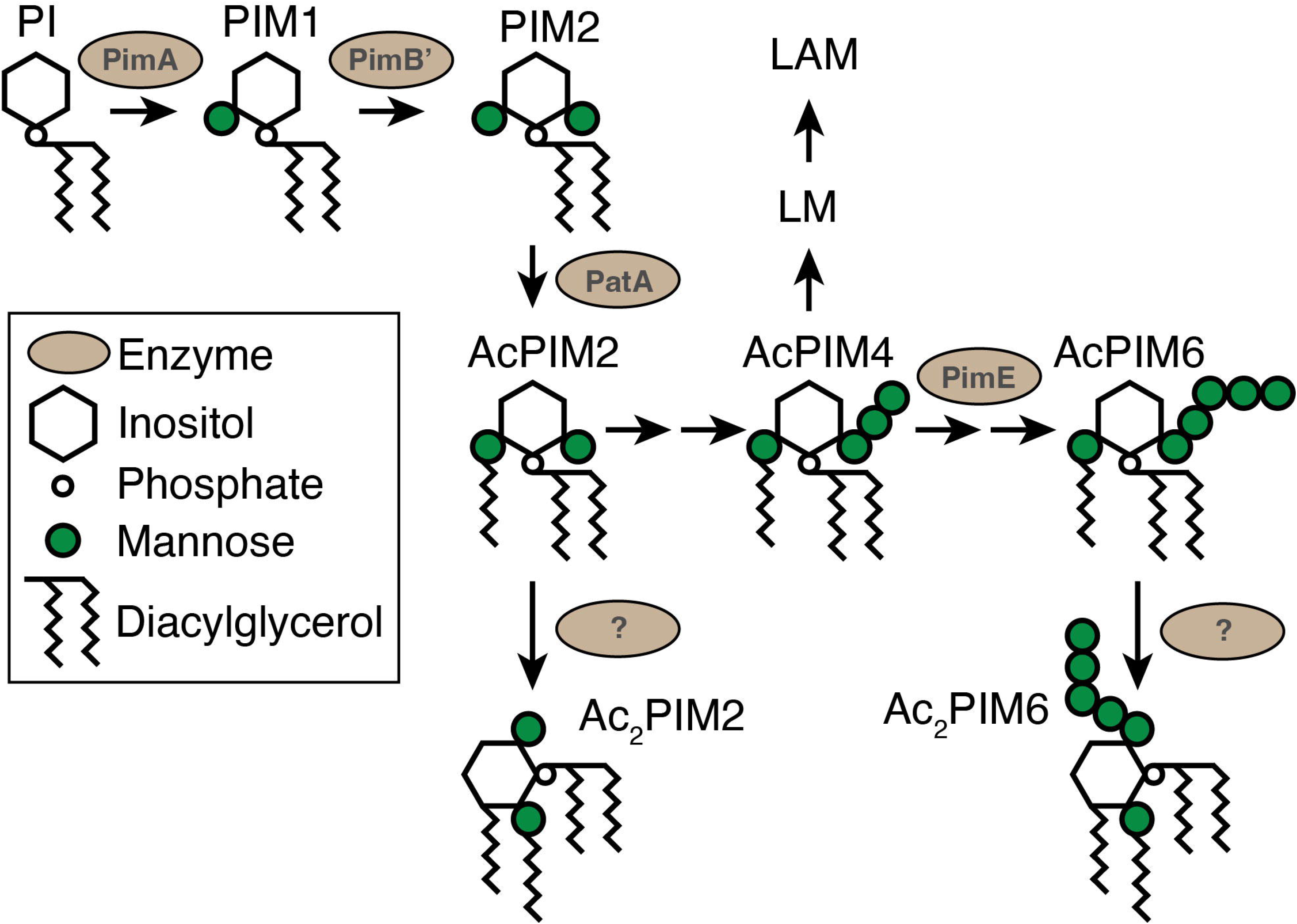
The PIM biosynthetic pathway. AcPIM2 and AcPIM6 are dominant PIM species while Ac_2_PIM2 and Ac_2_PIM6 are relatively minor secondary products. Other PIM species are considered as biosynthetic intermediates and may exist in lower quantities. Each step is mediated via a mannosyltransferase or an acyltransferase. While PimE mediates the fifth mannose transfer, the enzymes that mediate the third, fourth, and sixth mannose transfer has not been identified. The acyltransferase mediating conversion of monoacyl PIMs to diacyl PIMs has not been identified (indicated by question marks). Enzymes that mediate LM/LAM biosynthesis are not shown.

The roles of PIMs in mycobacterial physiology remain poorly characterized, but limited data suggest their importance as structural components of the plasma membrane. Genes encoding PimA and PimB’ are essential for *Mycobacterium smegmatis* under standard laboratory growth conditions (12, 13). *M. smegmatis ΔpatA* also shows severe growth defects (15). These early gene deletion studies indicate that one or more of the downstream products (*i.e*. PIMs / LM / LAM) is critical for growth. In contrast to the early-stage enzymes, growth defects of *M. smegmatis* Δ*pimE* are relatively mild, and dependent on media composition. When grown on Middlebrook 7H10 agar media, Δ*pimE* forms a small lumpy colony in contrast to a rugose and spread colony morphology of wildtype *M. smegmatis*. The growth defects are attributed to increased permeability of the Δ*pimE* cell envelope, making the concentration of copper in the medium toxic for the mutant (18). Electron microscopy revealed severe plasma membrane deformations in Δ*pimE* (16), suggesting that either the lack of AcPIM6 or the accumulation of AcPIM4 intermediate affects the plasma membrane integrity. In another study, an *M. smegmatis* mutant with a defective inositol monophosphate phosphatase accumulated less PIM2 and showed increased sensitivities to hydrophilic antibiotics (19). These studies are consistent with the idea that PIMs are important for plasma membrane integrity.

What is the significance of PIM inositol acylation? Under standard laboratory culture conditions, non-inositol-acylated PIMs (AcPIM2 and AcPIM6) are dominant, and inositol-acylated forms (Ac_2_PIM2 and Ac_2_PIM6) are quantitatively minor. Importantly, high salinity induces a gradual accumulation of inositol-acylated forms (20), suggesting that inositol acylation may be a stress response. Furthermore, in a recent study, we observed accumulation of Ac_2_PIM2 and Ac_2_PIM6 upon treatment with benzyl alcohol, a membrane fluidizer (21). In the current study, we examine this physiological response to benzyl alcohol in detail and demonstrate that it is a membrane stress response conserved across mycobacterial species.

## Materials and Methods

### Cell culture

*Mycobacterium smegmatis* (mc^2^155) (22) was grown in Middlebrook 7H9 medium supplemented with 15 mM NaCl, 0.2% (w/v) glucose, and 0.05% (v/v) Tween-80. All cultures were grown with shaking at 37°C except during heat treatments and short time course experiments. Colony forming units (CFU) were enumerated by spotting 5 μl of serially diluted cell cultures on Middlebrook 7H10 agar supplemented with 15 mM NaCl and 0.2% (w/v) glucose. To achieve the final working concentration of benzyl alcohol, 5 M benzyl alcohol in dimethyl sulfoxide (DMSO) was added to the culture. Heat was applied by placing the flasks in a pre-warmed water bath with occasional mixing.

For the short time course experiments, we concentrated the cell density as follows. Cells were grown to a log phase and centrifuged at 3,220x *g* for 10 minutes to obtain a pellet. The cell pellet was resuspended in 9 volumes of Middlebrook 7H9 medium. An aliquot (800 μl) of cells were then incubated in a glass tube in a prewarmed water bath at 37°C with benzyl alcohol.

Attenuated *Mycobacterium tuberculosis* H37Rv Δ*RD1* Δ*panCD* mc^2^6230 was grown in Middlebrook 7H9 supplemented with Middlebrook OADC supplement and 50 μg/ml pantothenic acid. *Mycobacterium abscessus* (ATCC19977) was grown in Middlebrook 7H9 supplemented with Middlebrook OADC supplement. *Corynebacterium glutamicum* (ATCC 13032) was grown in Brain Heart Infusion medium.

### Rapid heat-killing

To kill *M. smegmatis* cells rapidly without inducing heat-shock responses, 100 ml of cells in a 500-ml flask were microwaved for 30 seconds three times with brief mixing in between at 1,250 W.

### Lipid extraction

To harvest lipids, 25 OD_600_ units of cells were centrifuged at 3,220x *g* for 10 minutes and the pellet was resuspended in 20 volumes of chloroform/methanol (2:1, v/v), briefly vortexed, and sonicated. Following a 1.5-hour room temperature incubation, the suspension was spun down at 16,900x *g* on a microfuge for 1 minute and the supernatant was collected. This process was repeated with 10 volumes of chloroform/methanol (2:1, v/v), and then 10 volumes of chloroform/methanol/water (1:2:0.8, v/v/v) against the same pellet.

For the short time course experiments, 180 μL of cells were taken from the concentrated liquid cultures and mixed with 1.2 mL of chloroform/methanol (1:1, v/v) to obtain the final ratio of chloroform/methanol/water (10:10:3, v/v/v). Following a 1.5-hour incubation, the suspension was spun down at 16,900x *g* on a microfuge for 1 minute and the supernatant was collected. The pellet was then resuspended with 600 μL of chloroform/methanol (2:1, v/v) for additional lipid extraction.

For both methods, the combined extracts were dried under nitrogen stream, followed by an additional purification step using 1-butanol/water (2:1, v/v) phase partitioning as described previously (23). The butanol phase was dried and resuspended at 1 mg wet pellet equivalent per μL of water-saturated butanol.

### High-performance thin layer chromatography (HPTLC)

Purified glycolipids were developed on an HPTLC plate (silica gel 60, EMD Merck) in a solvent containing chloroform/methanol/13 M ammonia/1 M ammonium acetate/water (180:140:9:9:23, v/v/v/v/v). For glycopeptidolipids (GPLs) and trehalose dimycolates (TDM), lipid extracts were applied on an HPTLC plate and chromatographed with chloroform/methanol/water (9:1:0.1, v/v/v). Glycolipids were detected by orcinol staining. Phospholipids were detected by molybdenum blue staining.

We followed our published protocol to quantify PIMs (24). Briefly, 1 mM mannose was plated alongside lipid extracts at concentrations of 0.4, 0.6, 0.8, 1.0, 2.0, and 4.0 nmol to generate a standard curve. The stained bands were scanned and quantified in terms of mannose content using ImageJ visualization software.

### Antiseptic sensitivity assay

Cells were grown in 20 mL of Middlebrook 7H9 medium at 37°C to a log phase. After treating cells with benzyl alcohol for 5 or 60 minutes, the entire culture was spun down at 3,220x *g* for 10 minutes and then resuspended in the same volume of the medium. An aliquot (500 μl) of cell suspension was then incubated in a 2-mL microtube containing either benzethonium chloride (BTC) or sodium dodecyl sulfate (SDS) at 37°C with no shaking for 30 minutes. CFUs were enumerated as described above.

## Results

### Benzyl alcohol-induced inositol acylation is a biological and specific response

We first confirmed whether benzyl alcohol-induced inositol acylation is a biological phenomenon or an experimental artifact. Although it seemed unlikely, we were concerned that the addition of benzyl alcohol could influence lipid extraction efficiencies, *e.g*., benzyl alcohol may make the extraction of AcPIM2 and AcPIM6 less efficient, and that of Ac_2_PIM2 and Ac_2_PIM6 more efficient. To test, we first killed *M. smegmatis* by rapid heating and confirmed effective killing by CFU count (Fig. 2A). We then treated live and heat-killed cells with benzyl alcohol and analyzed extracted lipids by HPTLC. Rapid heating did not affect the PIM profile (Fig. 2B, compare lanes 1 and 3). Importantly, we found no accumulation of inositol-acylated PIMs from heat-killed cells that are treated with benzyl alcohol while Ac_2_PIM2 and Ac_2_PIM6 accumulated in live cells after a 60-minute benzyl alcohol treatment (Fig. 2B, compare lanes 2 and 4). In contrast to a nearly complete conversion of PIMs from inositol non-acylated to inositol acylated forms upon benzyl alcohol treatment (Fig. 2C), there were no major changes in other phospholipid species such as cardiolipin, phosphatidylethanolamine, and PI (Fig. 2D). Outer membrane lipids such as GPLs and TDM were also largely unaffected although there were quantitative changes in some lipid species (Fig. 2E). Together, these results indicate that inositol acylation of PIMs is a biological and robust modification of mycobacterial membrane elicited by benzyl alcohol.

**Figure 2.**
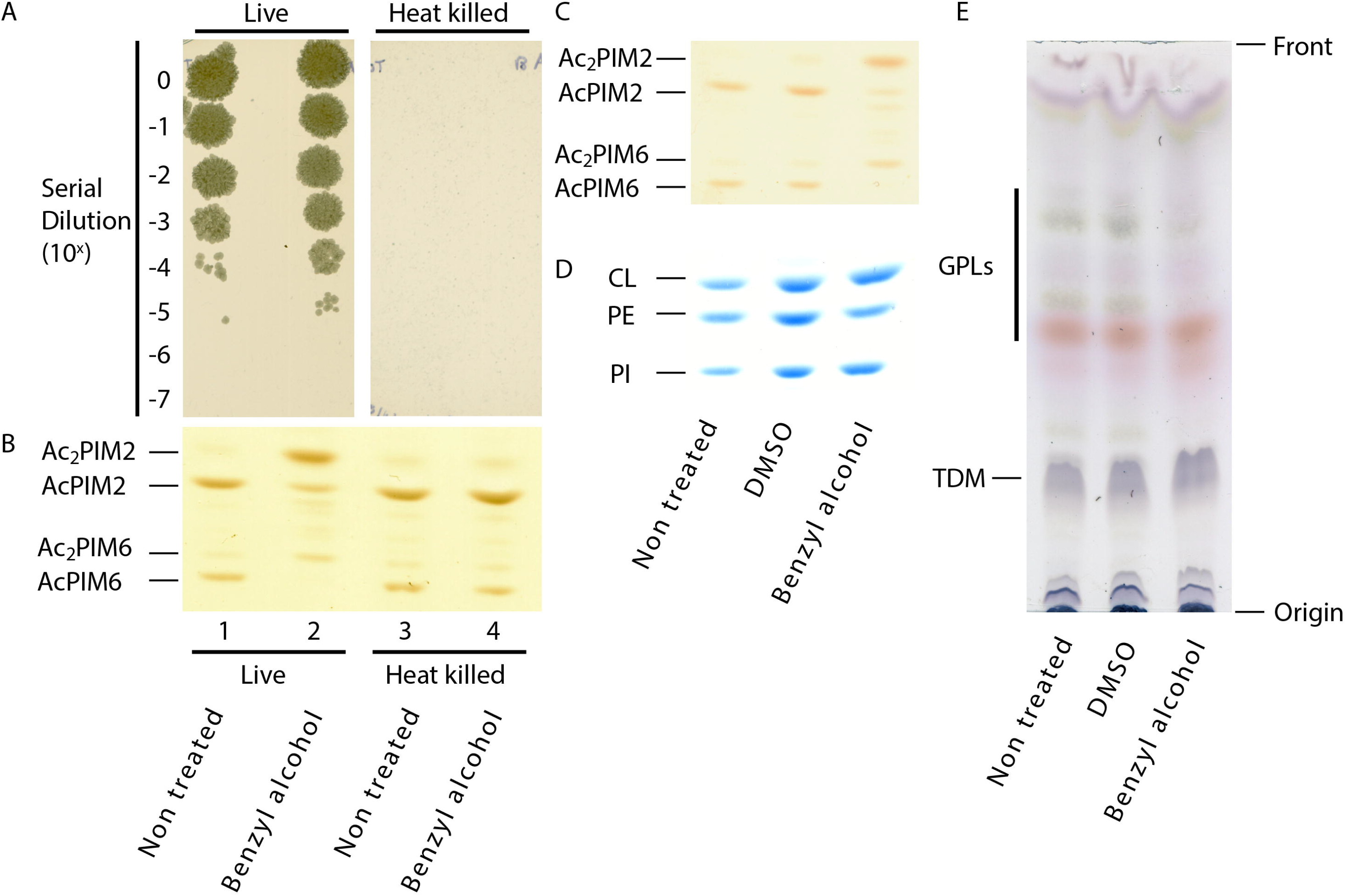
Effect of benzyl alcohol on *M. smegmatis* lipids. (A) Live and heat killed cells were treated with benzyl alcohol for 60 minutes, serially diluted, and plated onto Middlebrook 7H10 agar, confirming that rapid heating is effective in killing *M. smegmatis*. (B) HPTLC analysis of PIMs after heat killing followed by benzyl alcohol treatment. PIMs were chromatographed using a solvent containing chloroform, methanol, 13 M ammonia, 1 M ammonium acetate, and water (180:140:9:9:23, v/v/v/v/v) and visualized by orcinol staining. A region of the HPTLC plate corresponding to Rf of 0.22 - 0.48 is shown. (C-E) HPTLC analysis of lipids: (C) PIMs separated and visualized as in panel B (Rf = 0.20 – 0.48); (D) phospholipids separated as in panel B and visualized by molybdenum blue (Rf = 0.35 – 0.61); (E) GPLs and TDM separated by chloroform/methanol (9:1, v/v) and visualized by orcinol. DMSO, vehicle control. CL, cardiolipin; and PE, phosphatidylethanolamine. All experiments were repeated at least twice, and representative results are shown.

### Inositol acylation is induced by concentrations of benzyl alcohol that arrest cell growth

We used 100 mM benzyl alcohol in a previous study because it disrupts the membrane domain organization and arrests cell growth (21). However, we did not titrate the concentration of benzyl alcohol to examine if the concentration of benzyl alcohol correlates with PIM inositol acylation. When we tested lower concentrations of benzyl alcohol, we observed reduced levels of inositol acylation; complete PIM inositol acylation required 100 mM benzyl alcohol (Fig. 3A). To examine the effects of benzyl alcohol on growth, we monitored OD_600_ of *M. smegmatis* culture in the presence of benzyl alcohol (Fig. 3B). At 50 and 100 mM, benzyl alcohol was bacteriostatic, and there was little change in OD_600_ up to 24 hours. Cells treated with 25 mM benzyl alcohol grew at an intermediate rate between untreated and 50 mM benzyl alcohol. We examined the viability of cells up to 12 hours and found that there was only modest (less than one log) decline of CFU even after a 12-hour treatment with 100 mM benzyl alcohol (Fig. 3C). These data suggest that inositol acylation is a response to severe membrane stress that results in temporal arrest of the growth.

**Figure 3.**
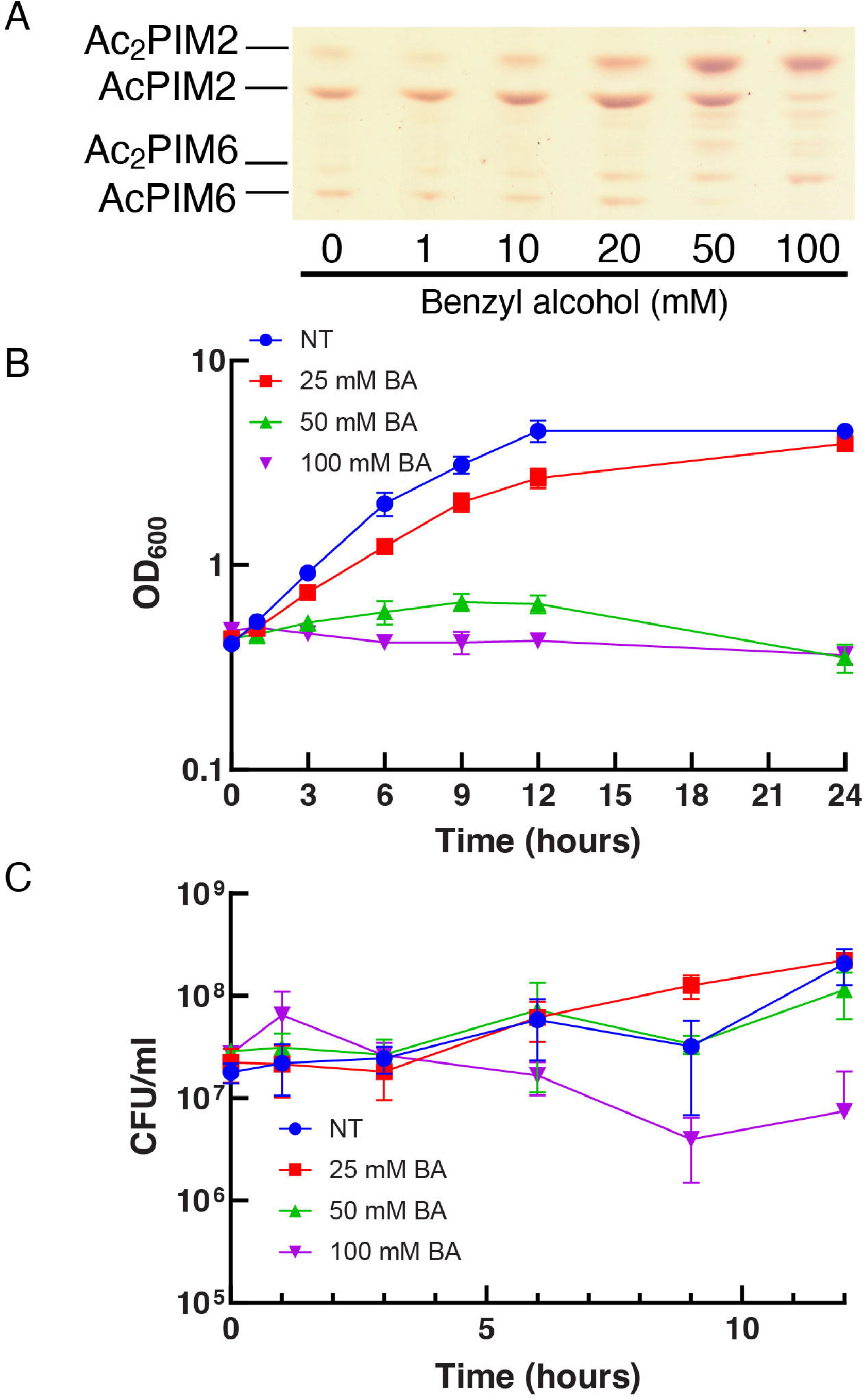
The extent of inositol acylation correlates with the concentration of benzyl alcohol. (A) HPTLC analysis of PIMs after treating with different concentrations of benzyl alcohol (Rf = 0.19 – 0.46). HPTLC was performed as in Fig. 2B. (B) Growth of *M. smegmatis* in the presence of increasing concentrations of benzyl alcohol. Experiments were performed in triplicate (average ± standard deviation). NT, non-treatment. BA, benzyl alcohol. (C) Viability of *M. smegmatis* during benzyl alcohol treatment. Experiments were performed in triplicate (average ± standard deviation).

### Inositol acylation is rapid

To examine the kinetics of the reaction, we took aliquots at different times after benzyl alcohol treatment and extracted lipids for HPTLC analysis. To take time points precisely in a short time course, we eliminated centrifugation steps to pellet cells because the extra time allowed the inositol acylation reaction to proceed before the reaction could be stopped by the addition of chloroform/methanol. To circumvent this problem, cells were treated with benzyl alcohol in a small volume at a cell density that is ~40 – 50 times denser than the normal culture cell density (see **Methods** for details). We then took a small aliquot at each time point over a course of 40 minutes, and directly suspended it in chloroform/methanol. At 100 mM benzyl alcohol, inositol acylation was detectable within 5 minutes, and nearly complete by 20 minutes (Fig. 4D). Similar, but less robust changes were observed at lower concentrations of benzyl alcohol (Fig. 4B and 4C). Without benzyl alcohol, there were only minor accumulations of diacyl species by 40 minutes (Fig. 4A), which were not observed under normal growth conditions (see Fig. 2). We suspect that the high cell density during the benzyl alcohol treatment may have elicited a minor stress response for unknown reasons. Taken together, these data further support that inositol acylation is a rapid biological response to benzyl alcohol.

**Figure 4.**
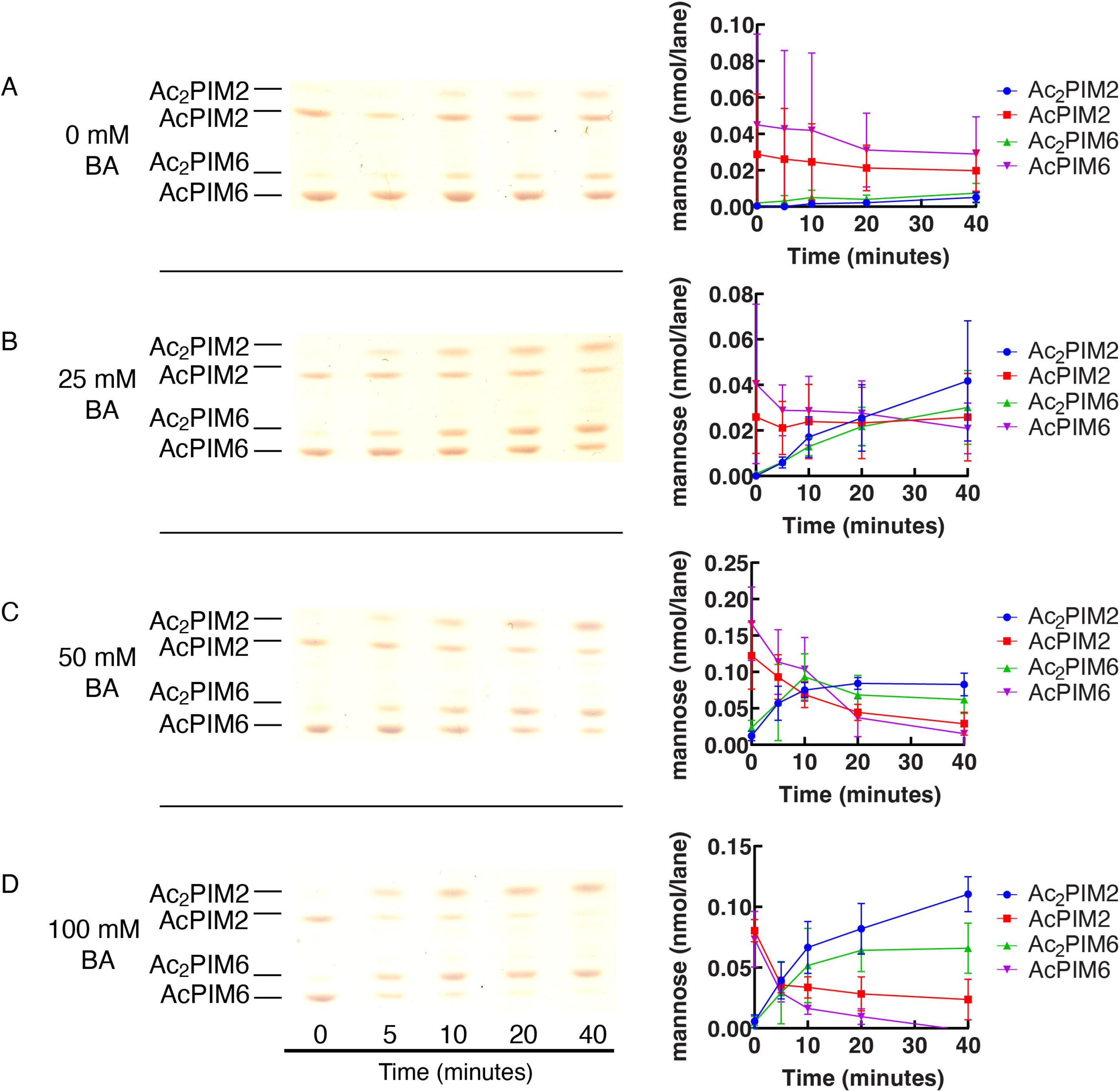
PIM acylation is rapid. (A-D) HPTLC of PIMs during a 40-minute treatment with (A) 0 mM (Rf = 0.18 – 0.45), (B) 25 mM (Rf = 0.18 – 0.45), (C) 50 mM (Rf = 0.21 – 0.47), and (D) 100 mM benzyl alcohol (Rf = 0.21 – 0.47). HPTLC was performed and PIMs were visualized as in Fig. 2B. BA, benzyl alcohol. Representative image of triplicate experiments is shown. For quantification, HPTLC plates were scanned, and band intensities of PIM species were quantified as mannose content (see **Methods** for details). Average ± standard deviation is shown in the graphs.

### Inositol acylation is independent of translation

Since inositol acylation was rapid and nearly complete in 20 minutes, it appeared likely that it is a direct enzymatic response without involving *de novo* protein synthesis. To test, we treated *M. smegmatis* cells with 10 μg/ml chloramphenicol for 10 minutes to inhibit protein synthesis followed by 100 mM benzyl alcohol to stimulate PIM acylation. Chloramphenicol is effective in inhibiting protein synthesis in *M. smegmatis* at this concentration (25), and was sufficient to arrest the cell growth (Fig. 5A). Treatment with chloramphenicol alone revealed no change in PIM profiles (Fig. 5B, compare lanes 1 and 3). When cells were treated with benzyl alcohol after chloramphenicol, we observed efficient inositol acylation (Fig. 5B, lanes 3 and 4), which was essentially identical to benzyl alcohol treatment without chloramphenicol addition (Fig. 5B, lanes 1 and 2). These data are consistent with the idea that benzyl alcohol-induced inositol acylation is a posttranslational event and does not require *de novo* protein synthesis.

**Figure 5.**
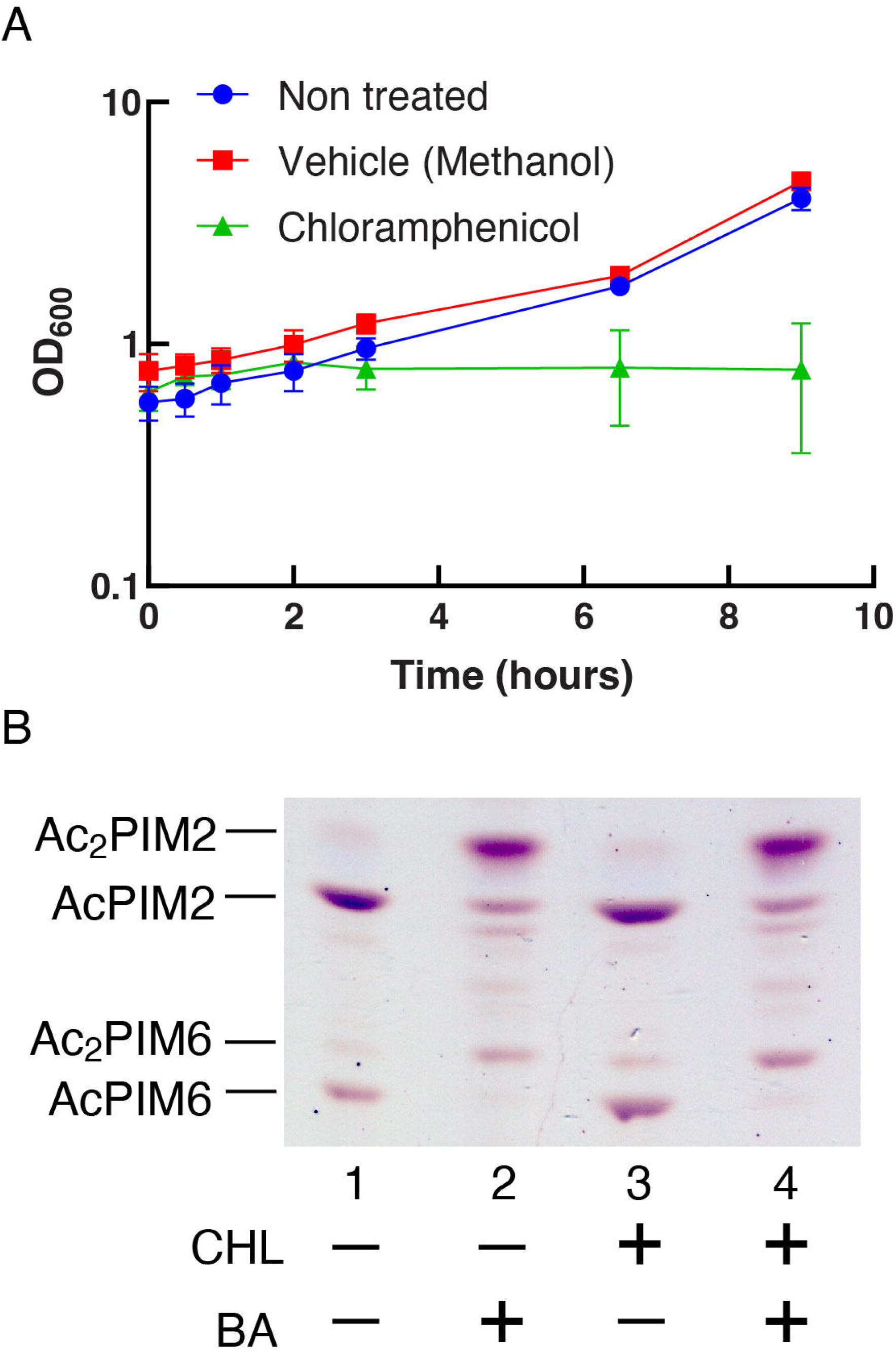
Inhibition of protein synthesis does not affect PIM inositol acylation. (A) Growth of *M. smegmatis* treated with 10 μg/ml chloramphenicol measured by OD_600_ over 24 hours. Experiments were performed in triplicate and average ± standard deviation is shown. (B) Cells were treated with 10 μg/ml chloramphenicol or methanol (vehicle control) for 10 minutes followed by 100 mM benzyl alcohol or DMSO (vehicle control) for 60 minutes. BA, benzyl alcohol. HPTLC analysis of PIMs is as described in Fig. 2B (Rf = 0.14 – 0.42). CHL, chloramphenicol. Experiment was repeated twice, and a representative result is shown.

### Inositol acylation is not immediately reversible

The data so far suggest that inositol acylation takes place rapidly when the membrane integrity is severely impacted. To examine if inositol acylation is a reversible reaction, we treated the cells with benzyl alcohol for 1 hour, spun down, and resuspended in fresh medium for recovery. The cells grew at a slower rate for the first ~4 hours after benzyl alcohol treatment (Fig. 6A). We took time points up to 6 hours of recovery and analyzed the PIM profiles by HPTLC. The levels of accumulated Ac_2_PIM2 and Ac_2_PIM6 declined over 40 minutes of the recovery period (Fig. 6B). Interestingly, AcPIM2 recovered its normal level in 10 minutes while AcPIM6 did not recover for the first one hour. Normal ratios of acyl/diacyl PIM species were not restored for 3 hours. These complex behaviors to restore the normal PIM profile imply that inositol acylation is not a simple reversible switch.

**Figure 6.**
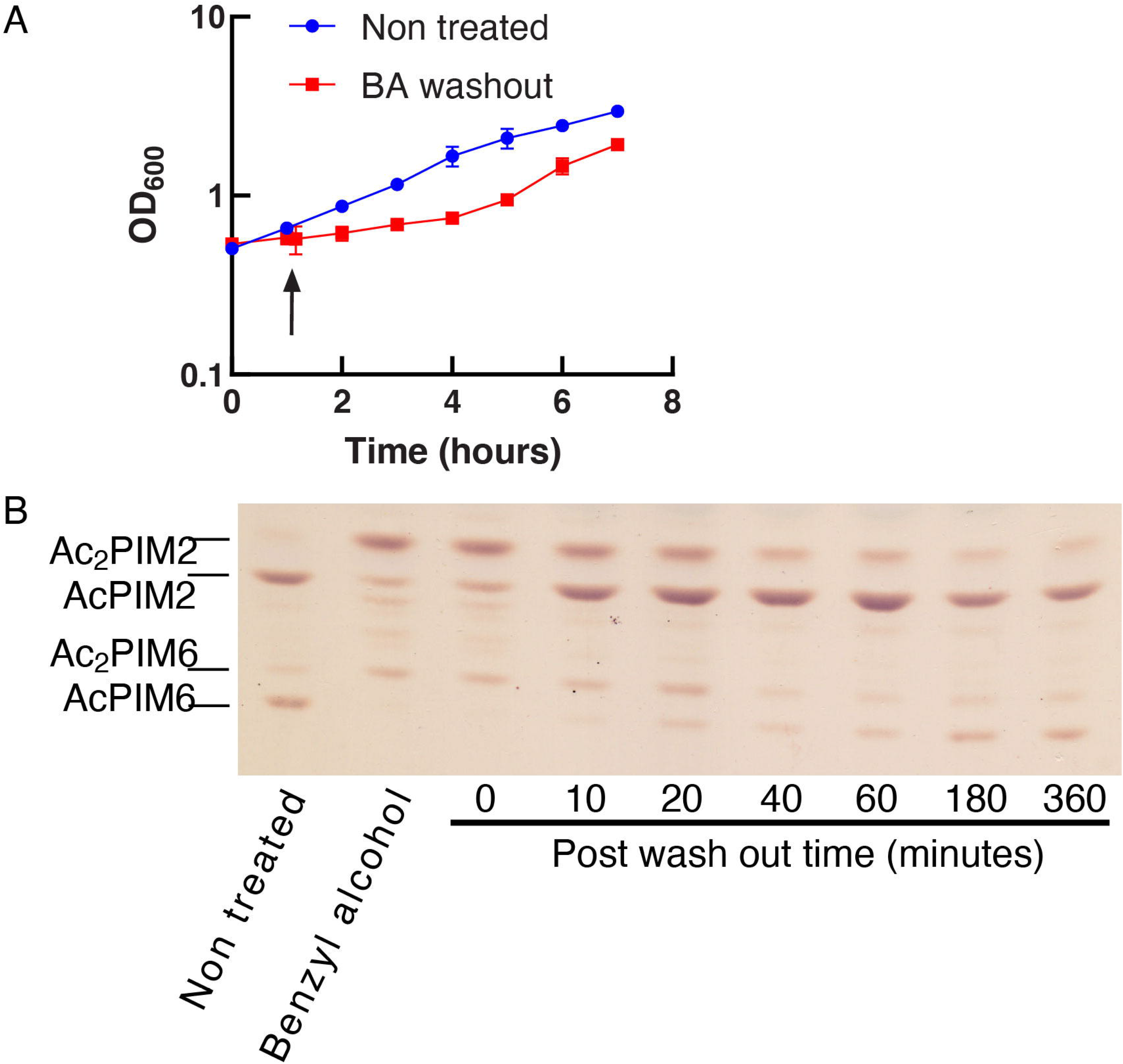
Recovery of PIM profile is gradual after washing out benzyl alcohol. (A) Growth of *M. smegmatis* during and after 60-minute benzyl alcohol treatment, monitored by OD_600_. Time 0 indicates the beginning of benzyl alcohol treatment. The black arrow indicates the 1-hour time point when benzyl alcohol was washed out. BA, benzyl alcohol. Experiments were performed in triplicate and average ± standard deviation is shown. (B) HPTLC analysis of PIMs after benzyl alcohol wash-out (Rf = 0.18 – 0.50). HPTLC was performed as in Fig. 2B. Experiment was repeated twice, and a representative result is shown.

### Inositol acylation is triggered by membrane fluidizing chemicals

Benzyl alcohol is a membrane-fluidizing chemical extensively used for both eukaryotic and prokaryotic membranes (26–36), and we have previously shown that benzyl alcohol disrupts membrane domain organization in *M. smegmatis* (21). To examine if the effect of benzyl alcohol on inositol acylation is due specifically to the chemical structure of benzyl alcohol or is a general property of aromatic alcohols, we tested different derivatives of benzyl alcohol. We tested 2-phenyl ethanol, 3-phenyl propanol, and 4-phenyl butanol, and found that all derivatives induced inositol acylation (Fig. 7A). Medium-length straight chain alcohols are also known to have membrane fluidizing properties (30, 36–40). To test their effect on inositol acylation, cells were treated with straight chain alcohols ranging from methanol to octanol. As with benzyl alcohol, these alcohols were tested at 100 mM except that hexanol and octanol were tested at 50 and 4 mM, respectively, because of their solubility limits in water. Butanol and longer chain alcohols all induced acylation to different extents, while short-chain alcohols such as methanol, ethanol, and propanol were ineffective (Fig. 7B).

**Figure 7.**
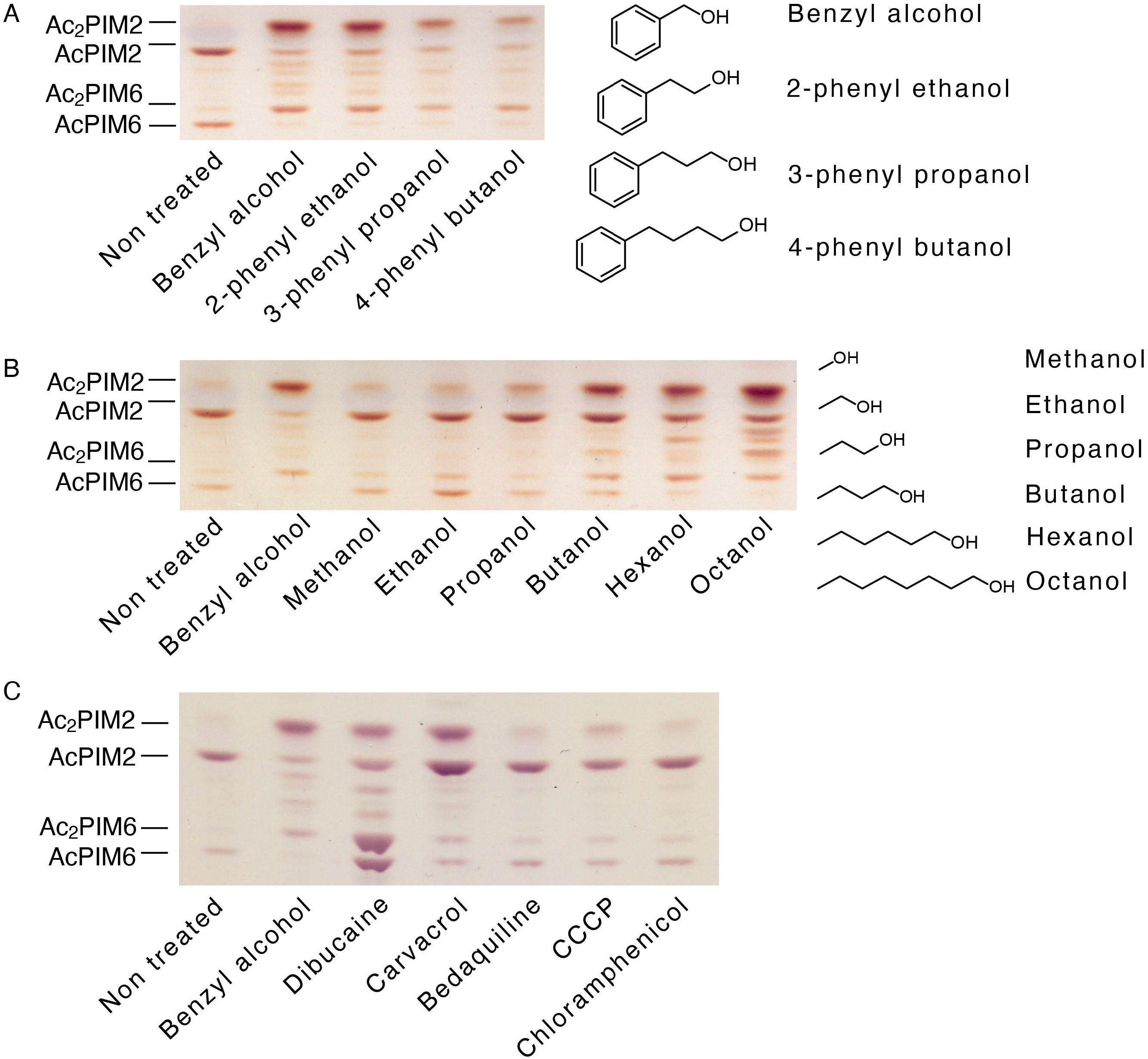
Inositol acylation in response to known membrane fluidizers. HPTLC analysis of PIMs extracted from *M. smegmatis* cells treated with (A) aromatic alcohols (Rf = 0.16 – 0.38), (B) straight chain alcohols (Rf = 0.17 – 0.39), or (C) other chemicals (Rf = 0.17 – 0.45). Cells were treated for 1 hour at 37°C with 100 mM alcohols (except hexanol and octanol at 50 mM and 4 mM, respectively), 600 μg/mL dibucaine, 1 mM carvacrol, 50 ng/ml bedaquiline, 25 μM CCCP, and 10 μg/ml chloramphenicol. Chloramphenicol was included as a control, which does not target the membrane (also see Fig. 5). HPTLC was performed as in Fig. 2B. All experiments were repeated at least twice, and representative results are shown.

Given these observations, changes in membrane fluidity appear to drive inositol acylation. To test further, we treated the cells with dibucaine, a local anesthetic known to fluidize membranes and disrupt liquid-ordered membrane domains (41–43). We have previously shown that dibucaine disrupts membrane organization in *M. smegmatis* (21). We found that dibucaine also induced inositol acylation (Fig. 7C) although the response was more complex than that of benzyl alcohol as AcPIM6 also increased in quantity in addition to the increased levels of Ac_2_PIM2 and Ac_2_PIM6. Numerous studies suggest that aromatic compounds in general can induce membrane fluidization (44–53). For example, carvacrol, a plant essential oil, induces changes in fatty acid compositions of bacterial membranes, which are indicative of membrane fluidization (54, 55). We tested the effect of carvacrol at 1 mM, a concentration approximately 5 times higher than the minimum inhibitory concentration (MIC) against *M. smegmatis* (56), and showed that carvacrol indeed was effective in inducing inositol acylation (Fig. 7C). Like dibucaine, we note the more complex responses to carvacrol, where the quantity of AcPIM2 was increased in concert with the increased levels of Ac_2_PIM2 and Ac_2_PIM6.

Bedaquiline is a tuberculosis drug, which targets the proton pump of ATP synthetase, leading to the disruption of redox balance (57, 58). It is an extremely lipophilic molecule with an estimated logP value of 7.74. We tested this drug at 50 ng/ml, approximately 5 times higher than the MIC against *M. smegmatis* (59). We also tested carbonyl cyanide m-chlorophenyl hydrazone (CCCP), a well-established uncoupler which collapses proton gradient in *M. smegmatis* (60, 61) and many other bacteria. These inhibitors did not affect the inositol acylation (Fig. 7C), arguing against the possibility that inositol acylation is a nonspecific response to disruption of any membrane functions.

### Benzyl alcohol-induced adaptation makes *M. smegmatis* more resistant to a membrane-active antiseptic

To examine the impact of benzyl alcohol-induced membrane remodeling in *M. smegmatis*, we tested the toxicity of antiseptic detergents, BTC and SDS, against cells that were pretreated with benzyl alcohol for either 5 or 60 minutes. After washing out benzyl alcohol by centrifugation, cells were challenged with BTC or SDS for 30 minutes. Cells were then serially diluted and plated on Middlebrook 7H10 agar for CFU determination. As shown in Fig. 8, cells from all conditions were killed to the same extent at 0.125% (w/v) SDS treatment, showing no effect of benzyl alcohol pretreatment to protect cells from the bactericidal effect of SDS. In contrast, cells pretreated with benzyl alcohol for 60 minutes became significantly more resistant to 25 μg/ml BTC, compared with untreated cells (Fig. 8). Although it was not statistically significant, even 5-minute benzyl alcohol treatment appeared to make the cells more resistant to BTC. These results imply that adaptation to benzyl alcohol exposure strengthens the integrity of the plasma membrane.

**Figure 8.**
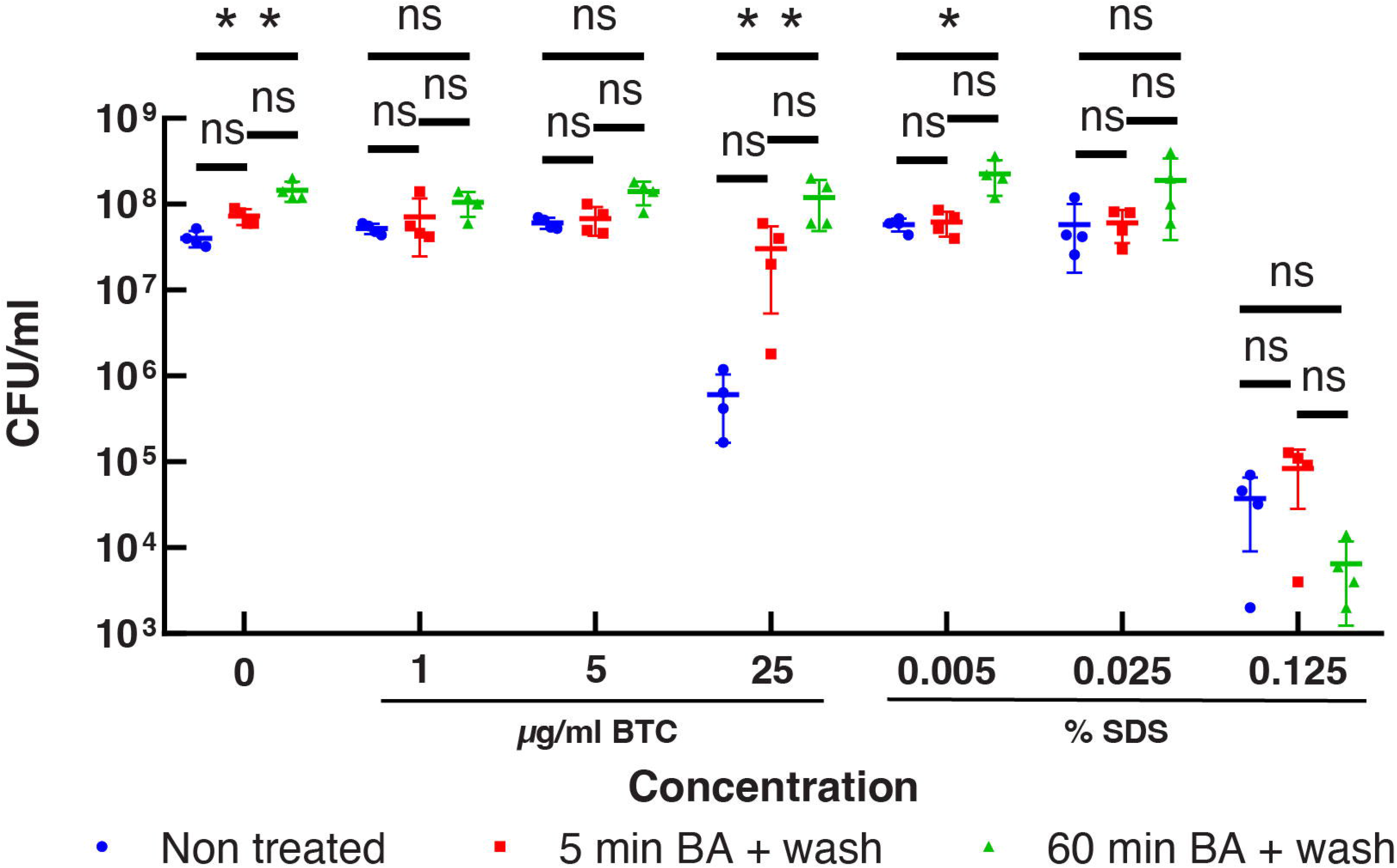
Benzyl alcohol pretreatment improves cell survival against an antiseptic detergent. Viability of *M. smegmatis* after treating with increasing concentrations of BTC or SDS. Prior to the detergent treatment, cells were subjected to no treatment (blue circle), 5-minute benzyl alcohol treatment and washout (red square), or 60-minute benzyl alcohol treatment and washout (green triangle). BA, benzyl alcohol. Experiment was done in quadruplicate. *, p = 0.0422. **, p =0.0049 (no detergent) or 0.0066 (25 μg/ml BTC) by Kruskal-Wallis test followed by Dunn’s multiple comparison test. In the absence of BTC or SDS challenges, the CFU showed a statistically significant increase upon treating cells with benzyl alcohol. We speculate that benzyl alcohol may help dispersing clumped cells, resulting in a small but consistent increase in CFU.

### Severe heat-shock also induces inositol acylation

Temperature is another factor that has been long known to influence plasma membrane composition and fluidity in bacteria (62–69), including mycobacteria (70). Therefore, we tested the effect of heat on inositol acylation. A relatively mild heat shock at 42°C induces the transcription of chaperonins such as GroES homolog Cpn10 and GroEL homologs Cpn60.1 and Cpn60.2 (71). However, when we treated cells at 42°C for 60 minutes, we observed only a mild increase of Ac_2_PIM2, and there were no apparent decreases of AcPIM2 and AcPIM6 (Fig. 9A). Consistent with a mild nature of 42°C heat shock, the cells started to regrow quickly without a substantial lag time (Fig. 9D). In contrast, when we treated cells at 50°C or 55°C, we observed substantial levels of accumulation of Ac_2_PIM2 and Ac_2_PIM6 (Fig. 9A). There were also concomitant decreases of AcPIM2 and AcPIM6. The CFU confirmed that the cells were not killed during the heat shock at 55°C (Fig. 9B), but the levels of the accumulated inositol-acylated PIM species remained unchanged even after 5 hours of recovery at 37°C (Fig. 9C). These observations suggest that there was nonlethal but severely growth-arresting damage to the cells at this temperature. Indeed, there was a much longer lag time (~12 hours) after the 55°C heat shock (Fig. 9D). These data are consistent with the data obtained using chemical fluidizers in that inositol acylation is a response to membrane fluidization and occurs when cells encounter severe, growth-arresting membrane damages.

**Figure 9.**
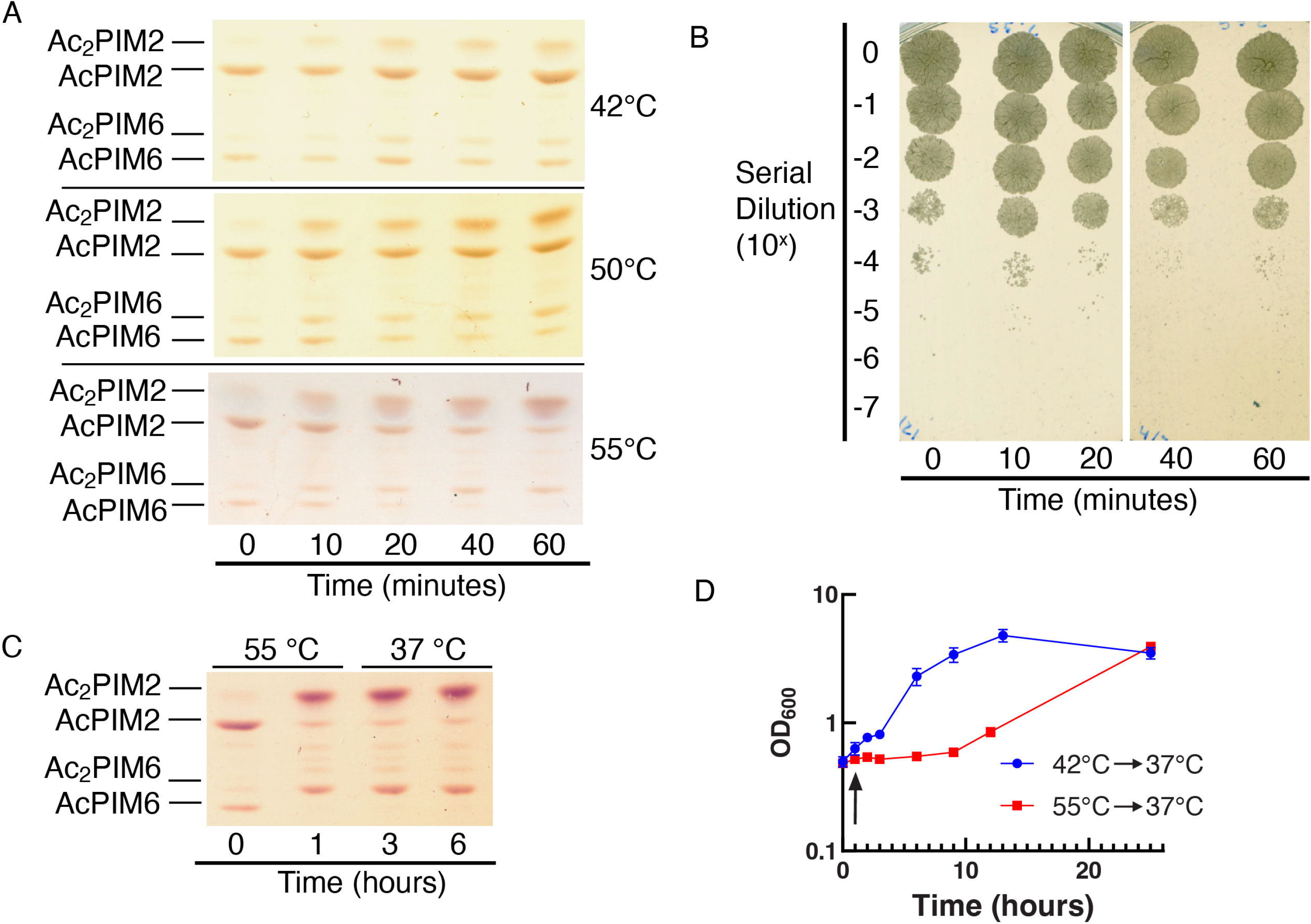
Heat shock induces PIM acylation. (A) HPTLC analysis of PIMs after heating cells at 42°C, 50°C, and 55°C (Rf = 0.18 – 0.45, 0.18 – 0.45, and 0.15 – 0.41, respectively). (B) Cells during the 55°C heat shock were serially diluted and plated onto Middlebrook 7H10 agar, confirming that cells remained viable at this temperature. (C) HPTLC analysis of PIMs from cells that were heat-shocked at 55°C for 1 hour and recovered at 37°C for 5 hours (Rf = 0.18 – 0.43). Data shown for panels A–C are representative results from two independent experiments. (D) Recovery after heat shock at 42°C (blue circle) or 55°C (red square). The black arrow indicates when heat shock was stopped (at 1 hour). The experiment was done in triplicate.

### Inositol acylation is evolutionarily conserved

To determine if PIM acylation is evolutionarily conserved, we treated *Mycobacterium tuberculosis*, *Mycobacterium abscessus*, and *Corynebacterium glutamicum* with 100 mM benzyl alcohol. In contrast to the fast-growing and nonpathogenic *M. smegmatis*, *M. tuberculosis* and *M. abscessus* are slow- and fast-growing pathogens, respectively. Given the distinct lineages of these *Mycobacterium* species, there were clear differences in PIM profiles among them. In particular, the HPTLC mobilities (Rf values) of Ac_2_PIM2 and Ac_2_PIM6 in *M. abscessus* were different from those of *M. smegmatis* and *M. tuberculosis* (Fig. 10), suggesting different fatty acid compositions. Nevertheless, *M. tuberculosis* and *M. abscessus* produced acylated PIMs in response to benzyl alcohol in a manner similar to *M. smegmatis* (Fig.10). In contrast, *C. glutamicum*, which produces AcPIM2 and not AcPIM6 (72), did not acylate AcPIM2 (Fig. 10A), indicating that inositol acylation is a conserved response in the genus *Mycobacterium*.

**Figure 10.**
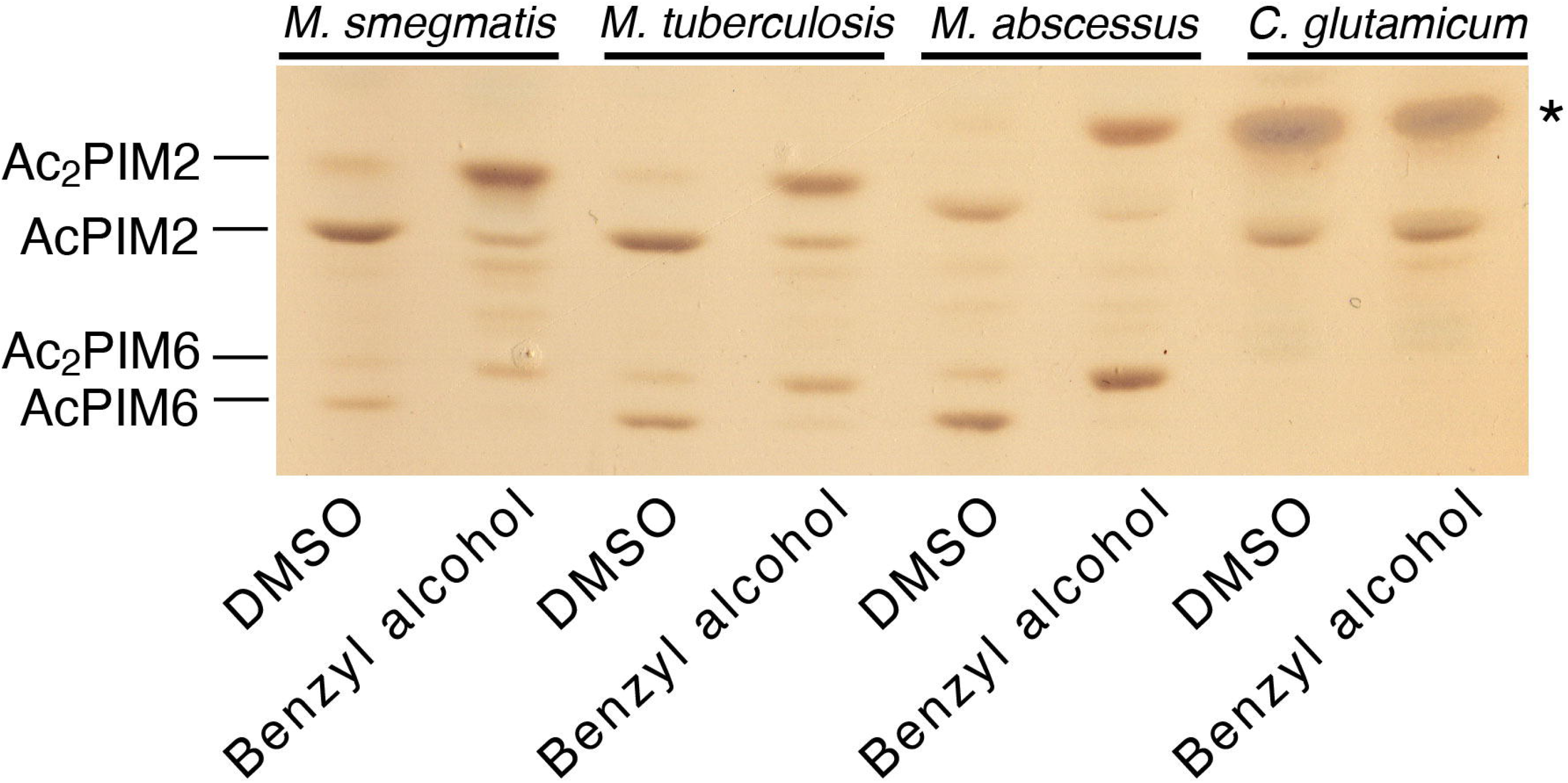
Benzyl alcohol treatment of other *Mycobacterium* and *Corynebacterium* species. HPTLC of lipids from *M. smegmatis, M. tuberculosis, M. abscessus*, and *C. glutamicum* treated with 100 mM benzyl alcohol for 1 hour (Rf = 0.16 – 0.45). HPTLC was performed as in Fig. 2B. Note that *C. glutamicum* does not produce PIM6 species. *, trehalose monocorynomycolate.

## Discussion

In this study, we demonstrated that PIM inositol acylation is a stress response to severe bacteriostatic membrane fluidization. Alteration of membrane lipid composition can be achieved through three distinct processes: 1) component biosynthesis (*e.g*. fatty acid and acyl-CoA); 2) lipid assembly (*e.g*. phospholipid biosynthesis); and 3) lipid modification and catabolism (73). Inositol acylation is well known as a modification of eukaryotic GPI anchors from human to protozoan parasites (74, 75). However, it primarily functions as a checkpoint in the GPI precursor biosynthesis, where acyl modification of inositol, mediated by the acyltransferase PIG-W in the endoplasmic reticulum, is often removed by the inositol deacylase PGAP1 prior to trafficking to the Golgi apparatus (76–79). Therefore, eukaryotic inositol acylation is an enzymatic step in the lipid assembly process. In contrast, mycobacterial PIM inositol acylation is a modification of mature lipid molecules. There are several known examples of lipid modification and catabolism. Among them, direct desaturation or deacylation/reacylation of membrane lipids for cold acclimatization is well characterized in both eukaryotes and bacteria (73, 80–82). However, these direct and rapid modifications of lipids still take hours, and often involve transcriptional upregulations (82–86). This is in stark contrast to PIM inositol acylation, which is nearly complete in 20 minutes. The only other example, to our knowledge, is the unsaturation of cyclopropane fatty acyl moieties of phospholipids in response to toluene in *Pseudomonas putida*, which was also a rapid mass reaction complete in about 20 minutes (87). Thus, PIM inositol acylation represents one of the fastest mass conversions of mature lipid species in nature.

What is the physiological significance of inositol acylation? Multi-acylated forms of PIMs have been known since the 1960s (88–91), and the inositol acylation was formally demonstrated in 1999 (92). While PIMs are suggested to be major components of the plasma membrane and play roles in maintaining plasma membrane integrity (6, 16), specific roles of inositol acylation remained obscure. Our study demonstrated that inositol acylation takes place rapidly in response to bacteriostatic membrane fluidization stress and is conserved in pathogenic species such as *M. tuberculosis* and *M. abscessus*. In contrast, inositol deacylation is a slower process, suggesting that inositol acylation may be a last resort defense against severe membrane damages rather than a reversible switch to finetune membrane fluidity. Exposures to high salt concentrations led to an accumulation of Ac_2_PIM2 in *M. tuberculosis* (20). This response was a slower response over several days and inositol acylation was incomplete even after a three-day incubation. Therefore, it is distinct from the rapid and complete conversion of PIM species reported in this study. It would be interesting to examine if salinity affects membrane fluidity or induces PIM inositol acylation independent of sensing membrane fluidity changes.

Notably, benzyl alcohol treatment not only induces inositol acylation but also makes *M. smegmatis* more resistant to BTC, a quaternary ammonium surfactant. We speculate that inositol acylation fortifies the plasma membrane by increasing its rigidity. However, we acknowledge that inositol acylation is not the only response to membrane fluidization (93–95), and the precise physiological significance of inositol acylation remains to be defined. Both benzyl alcohol and heat can exert other effects on the membrane and other cellular structures. For example, we found that heat treatment at 55°C was substantially more taxing for the cell than 100 mM benzyl alcohol treatment, resulting in much slower recovery process. Such a high temperature treatment may denature many more proteins than benzyl alcohol treatment, likely activating heat shock responses to restore protein homeostasis. Interestingly, heat shock proteins not only act as protein chaperones, but also bind membrane and protect its integrity (96). In *M. smegmatis*, when the chaperone DnaK is depleted, the cells become more heat-sensitive and membrane-permeable (25), suggesting a potential role for DnaK in membrane protection against heat shock. Furthermore, benzyl alcohol can induce heat-shock responses in other organisms in the absence of heat (31, 97, 98). These observations illuminate complex responses to membrane-fluidizing treatments and highlight the importance of determining precise contributions of inositol acylation to maintaining membrane fluidity and protecting membrane integrity. Further studies are needed to define the molecular mechanisms that govern the membrane stress response uncovered in this study. Identifying the inositol acyltransferase or deacylase is a critical future direction that will allow us to further examine the regulatory mechanism and physiological significance of inositol acylation.

PIMs are proposed to be enriched in the plasma membrane and contribute to a less fluid and less permeable plasma membrane (6). The enrichment of PIMs in the plasma membrane implies that the enzyme that mediates inositol acylation is also associated with the plasma membrane. Since the active site of the mannosyltransferases involved in the synthesis of polar PIMs (*e.g*. PimE) is suggested to be on the periplasmic side of the plasma membrane (16, 23), it seems likely that AcPIM6 is produced in the outer leaflet of the plasma membrane. Rapid conversion of AcPIM6 to Ac_2_PIM6 suggests that the enzyme that mediates inositol acylation is also on the outer leaflet of the plasma membrane. Assuming that the same enzyme mediates the inositol acylation of both AcPIM2 and AcPIM6, we speculate that AcPIM2 is likely available on the outer leaflet of the plasma membrane as well. PatA is an acyl-CoA-dependent acyltransferase that mediates the acylation step to produce AcPIM2 and is associated with the inner leaflet of the plasma membrane (15). Since we predict that the inositol acyltransferase is on the periplasmic side where acyl-CoA may not be readily available, we further speculate that this enzyme may be evolutionarily distinct from PatA.

We have previously shown that membrane domain organization is disrupted by benzyl alcohol and dibucaine (21). However, it remained largely unknown if mycobacteria can sense and respond to plasma membrane fluidity changes. Our current study demonstrated for the first time that mycobacteria possess a mechanism to respond to plasma membrane fluidization. How mycobacteria sense membrane fluidity is an open question. The mycobacterial cell envelope is 100 - 1,000 times less permeable to hydrophilic molecules than that of *E. coli* (99, 100), and the impermeability has been attributed to the number and property of porins in the outer membrane and low fluidity of outer membrane lipids. However, plasma membrane fluidity can be another major factor that contributes to overall cell envelope permeability (101). Our study revealed a rapid activation of inositol acyltransferase reaction, potentially suggesting direct sensing of membrane fluidity changes by the enzyme itself. Alternatively, it is possible that membrane fluidity changes are sensed by a membrane-embedded sensor that activates a signaling cascade, which includes the activation of the inositol acyltransferase. The Phage shock protein (Psp) response system senses membrane damage and repairs proton leakage across plasma membrane (102). CCCP activates the Psp system in mycobacteria. Since CCCP had no effect on inositol acylation, it is unlikely that the Psp system is involved in the activation of the inositol acyltransferase. DesK/R is one of the two-component regulatory systems in *Bacillus subtilis* (103). The sensor kinase DesK measures the membrane thickness, which thickens as temperature decreases (103–105). Upon activation, the cognate response regulator DesR activates the transcription of genes important for cold adaptation, among them being the fatty acid desaturase DesA, which facilitates the synthesis of unsaturated fatty acids to maintain the membrane fluidity at low temperature (103, 106). A similar response mechanism may be present in mycobacteria to monitor the membrane fluidity. However, since inositol acylation was not affected by the inhibition of translation by chloramphenicol, it may not be through a typical two-component response regulator. A more direct mechanism must exist, but the molecular mechanism remains unknown.

## Data Availability

All data are contained within the manuscript.

## Acknowledgements

*M. tuberculosis mc^2^6230 ΔRD1/panCD* strain was a kind gift from Dr. William R. Jacobs Jr. (Albert Einstein College of Medicine). We thank Casey Albano and Alkmini Diamantidi for help with experiments.

## Abbreviations

BTC: benzethonium chloride
CCCP: carbonyl cyanide m-chlorophenyl hydrazone
CFU: colony forming unit
DMSO: dimethyl sulfoxide
GPI: glycosyl phosphatidylinositol
GPL: glycopeptidolipid
HPTLC: high-performance thin later chromatography
LAM: lipoarabinomannan
LM: lipomannan
MIC: minimum inhibitory concentration
PI: phosphatidylinositol
PIM: phosphatidylinositol mannoside
SDS: sodium dodecyl sulfate
TDM: trehalose dimycolate

